# Properties of the epigenetic clock and age acceleration

**DOI:** 10.1101/363143

**Authors:** Louis El Khoury, Tyler Gorrie-Stone, Melissa Smart, Amanda Hughes, Yanchun Bao, Alexandria Andrayas, Joe Burrage, Eilis Hannon, Meena Kumari, Jonathan Mill, Leonard C Schalkwyk

## Abstract

**Background:** The methylation status of numerous CpG sites in the human genome varies with age. The Horvath epigenetic clock used a wide variety of published DNA methylation data to produce an age prediction that has been widely used to, predict age in unknown samples, and draw conclusions about speed of ageing in various tissues, environments, and diseases. Despite its utility, there are a number of assumptions in the model that require examination. We explore the characteristics of the model in whole blood and multiple brain regions from older people, who are not well represented in the original training data, and in blood from a cross-sectional population study.

**Results:** We find that the model systematically underestimates age in tissues from older people. A decrease in slope of the predicted ages were observed at approximately 60 years, indicating that some loci in the model may change differently with age, and that age acceleration measures will themselves be age-dependent. This is seen most strongly in the cerebellum but is also present in other examined tissues, and is consistently observed in multiple datasets. An apparent association of Alzheimer’s disease with age acceleration disappears when age is used as a covariate. Association tests in the literature use a variety of methods for calculating age acceleration and often do not use age as a covariate. This is a potential cause of misleading findings.

**Conclusions:** Associations of phenotypes with age acceleration should be evaluated cautiously, and chronological age should be included as a covariate in all analyses.

## Background

Cellular differentiation and growth are orchestrated by epigenetic modifications. DNA methylation is the most stable and easily assayed epigenetic mark and it is indicative of many changes that happen throughout life. Different cell types have dramatically different DNA methylation profiles and there is a substantial literature on CpG loci whose DNA methylation status is associated with participant age in cross sectional studies (Hannum *et al.*, 2013; Horvath, 2013; Spiers *et al.*, 2015; Zbieć-Piekarska *et al.*, 2015). These sites thus appear to change over time, which, amongst other processes, could reflect developmental changes, cumulative environmental effects, and changes in cell-type composition. Identifying these sources of variation could give insights into multiple age-related processes, and also provide a way of estimating the age of study participants at the time of sample collection. There are changes to the epigenome with age, the majority of studies have shown global demethylation (Wilson *et al.*, 2007; Bjornsson *et al.*, 2008; Zampieri *et al.*, 2015), although there is hypermethylation in CpG islands within gene promoters (Bell *et al.*, 2012).

Horvath (2013) used a large collection (n > 8000) of publically available Illumina HumanMethylation array data on multiple tissue types to train and test a model for age prediction from 353 CpG loci. This ‘epigenetic clock’ is extremely valuable as a way of estimating study participant ages, and possibly as an indicator of whether there are alterations in the ageing rate of certain tissues or individuals. With this in mind, it is attractive to try to relate these epigenetic changes with age related conditions such as dementia and Alzheimer disease (AD) (Zampieri *et al.*, 2015), frailty (Breitling *et al.*, 2016) and all-cause mortality (Marioni *et al.*, 2015). However, as the model was derived using data collected at a single time point and does not include within person changes it therefore does not directly assess ageing.

Ageing is a complex process described as a slow and gradual decline in physiological integrity leading to diminished function (López-Otín *et al.*, 2013). Many factors such as genomic instability, epigenetic alterations, mitochondrial dysfunctions and others, have been described by Lopez-Otin *et al.* (2013) as the hallmarks of ageing. Although these changes are involved with ageing and longevity, it is still unclear how these modifications are affected by the cells’ and tissues’ interaction with the environment. Although the epigenetic clock developed by Horvath (2013) provides an estimate of age, the testing data used in generating this clock did not have a large representation of brain tissue from elderly individuals and as such it is unclear if the clock is accurate in older age groups, or those with age-related diseases.

We have previously published an epigenome-wide association study (EWAS) in AD, utilizing four brain tissues and pre-mortem blood, and demonstrated DNA methyation differences at specific loci in a tissue-specific manner (Lunnon *et al.*, 2014). This dataset offers an ideal opportunity to investigate the properties of the Horvath (2013) model on different tissues in both elderly non-demented individuals and AD sufferers. We further explore the properties of the model using a cross sectional population sample from the *Understanding Society* study, which has a wide range of ages.

Because the AD effects found in Lunnon *et al* (2014) and De Jager *et al* (2014) are quite small in magnitude and in a set of probes not overlapping with the ones used by the Horvath (2013) model, we did not expect to see ‘age acceleration’ associated with AD. For there to be an association the loci included in the model would be exhibiting a subtle but pervasive AD effect, too small to be identifiable on a probewise analysis, but perhaps reflective of a common variable-rate biological ageing clock mechanism. Age acceleration associated with disease states or environmental factors might support the notion such a mechanism does exist, but we would like to caution that in a field where many variables are being tested against age acceleration (which can also be defined in several ways) one would expect a number of positive results by chance.

## Results

We first used the Horvath Epigenetic Clock to predict the DNAm ages of participants from two case-control studies of Alzheimer Disease (AD) included in our original publication, 390 brain and blood samples from 92 individuals in the London cohort (cohort 1) (Lunnon *et al.*, 2014) and 280 brain samples from 147 individuals in the Mount Sinai cohort (cohort 2) (Smith *et al.*, 2018). As shown in table 1, in these elderly individuals DNAm ages are consistently underestimated. As can be seen from figure 1, DNAm ages clearly correlated with actual ages, but strikingly for all tissues of the London cohort the slope is less than 1 (figure 1 A-F, dotted line).

**Figure 1.**
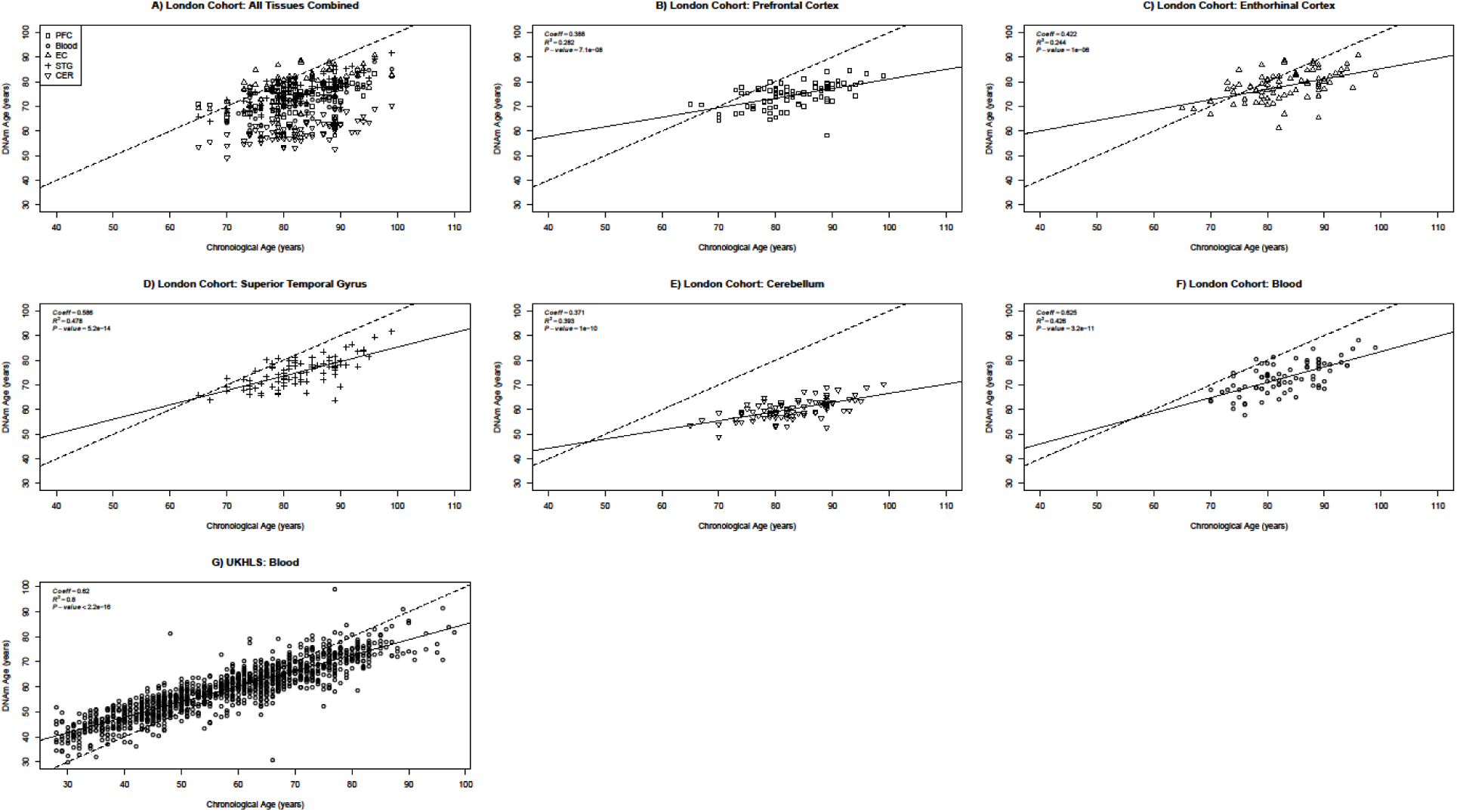
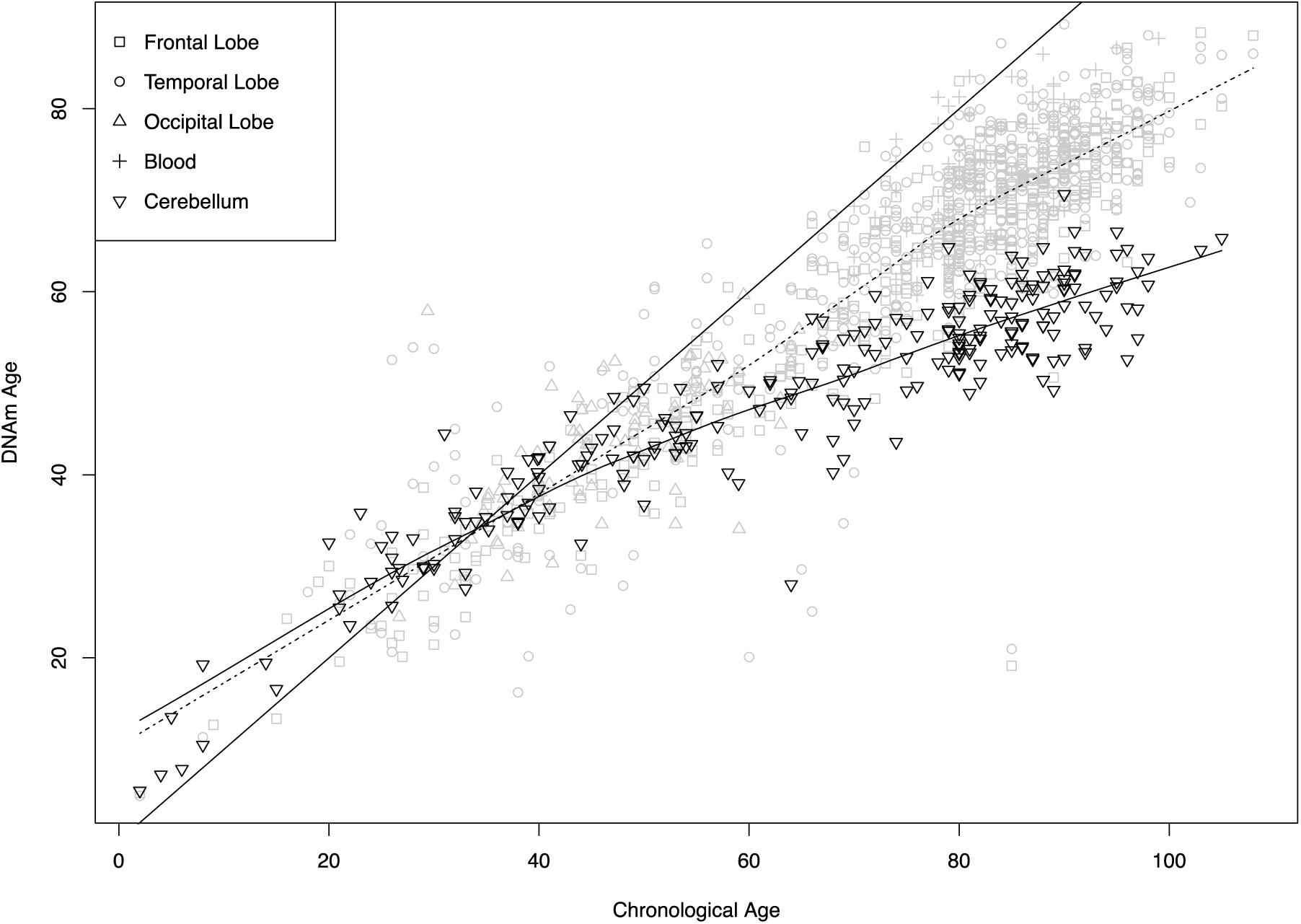
Scatterplot of Chronological vs DNAm ages of brain and blood samples. Each point corresponds to an independent sample. The dotted line is the y=x bisector line and the solid lines correspond to the regression line of each tissue. PFC: prefrontal cortex, EC: enthorhinal cortex, STG: the superior temporal gyrus, CER: cerebellum (data from Lunnon *et al* 2014 for panels A-F and Gorrie-Stone *et al* (2018, under peer-review) for panel G).

**Table 1.**
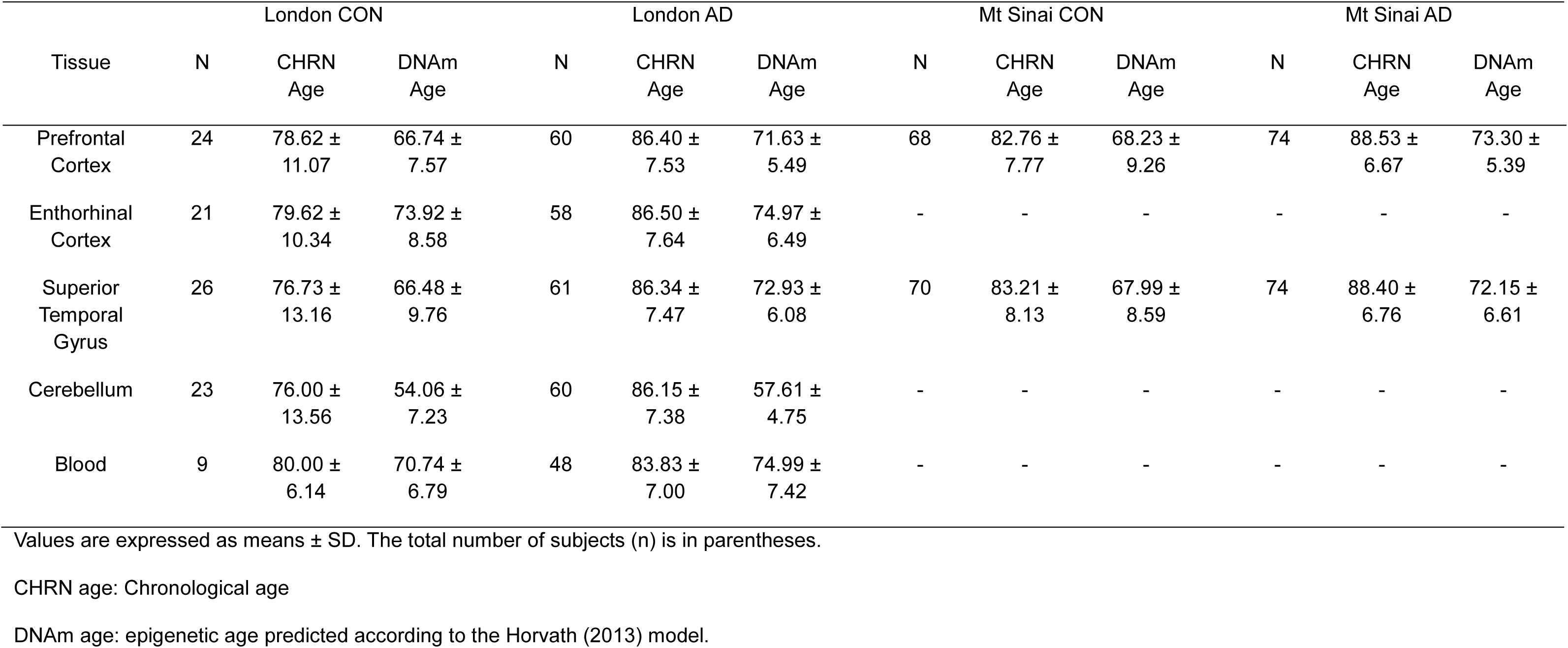
Chronological ages of samples obtained from the London Cohort

This picture is further confirmed by the data from the UKHLS population study. Here blood samples are obtained from participants across the adult age span and confirm the observation that DNAm ages are increasingly underestimated with advanced age (figure 1G). In figure 2 we show that the same effect holds across a variety of brain tissue data sets listed in table 3. In particular, the cerebellum is severely under predicted; far more than other brain and blood tissues (figure 2). Our consistent findings in tissue samples, from patients and a population study of healthy participants indicates that the results are robust and not specific to population or sample characteristics.

**Figure 2.**
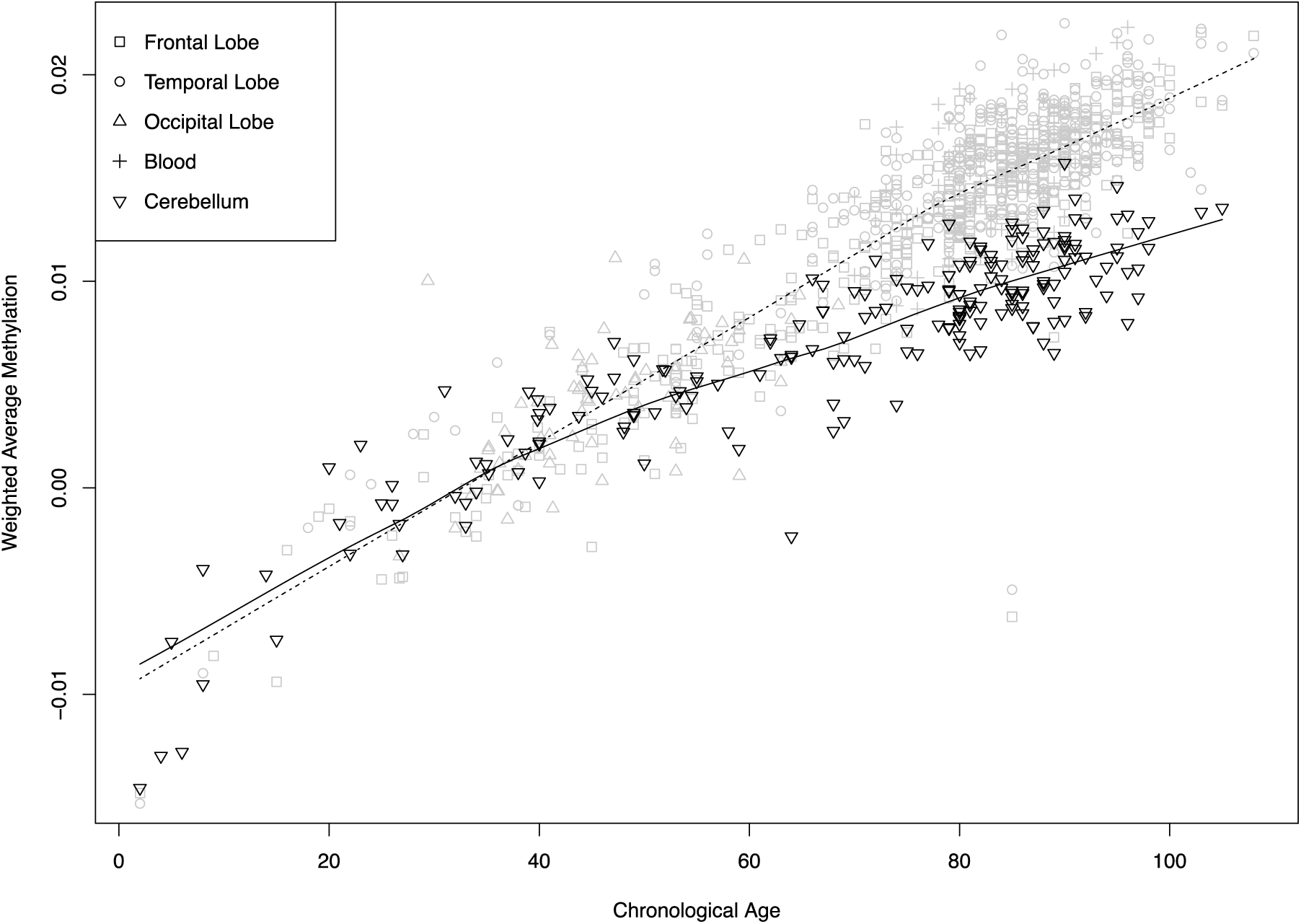
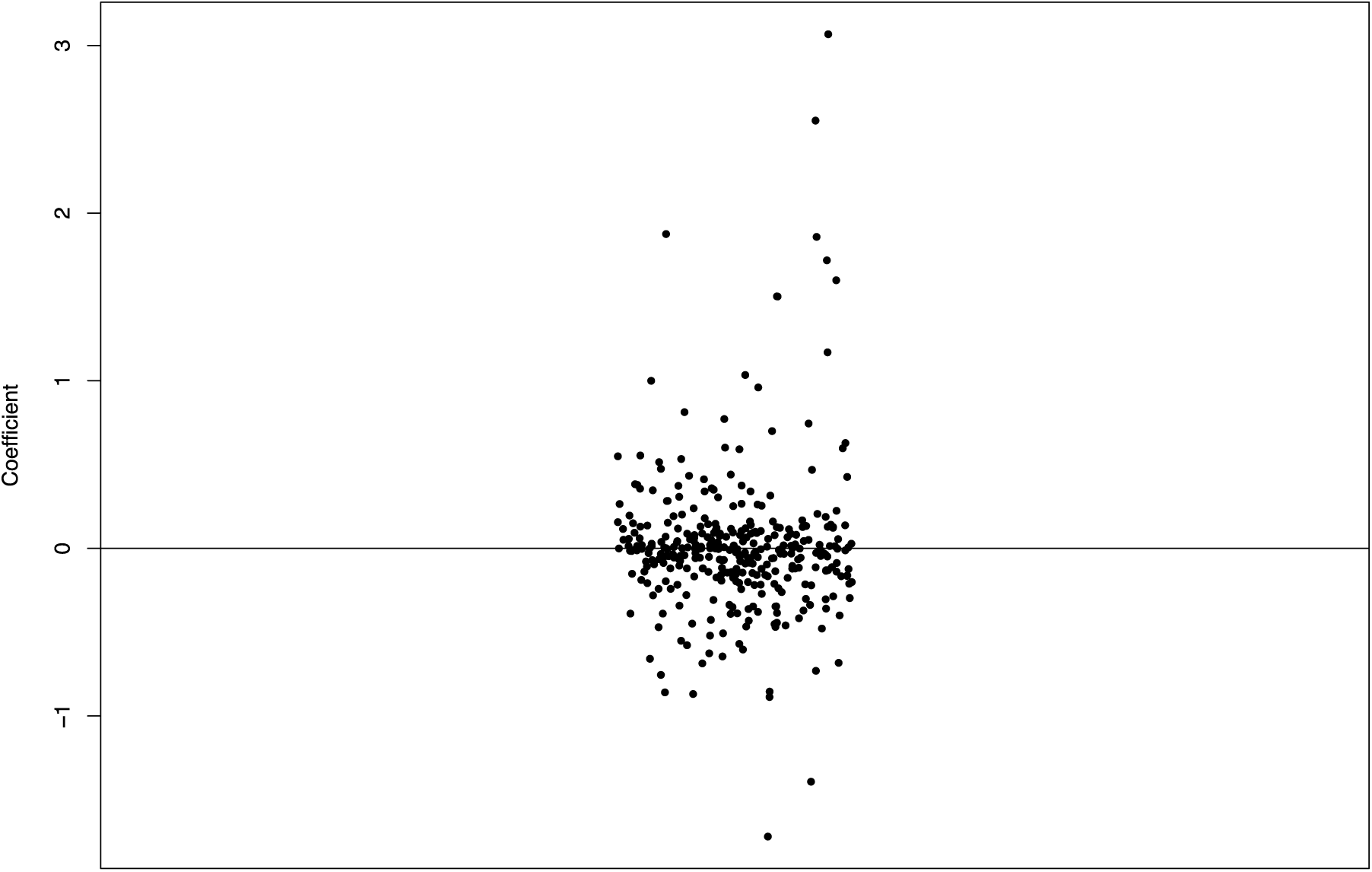
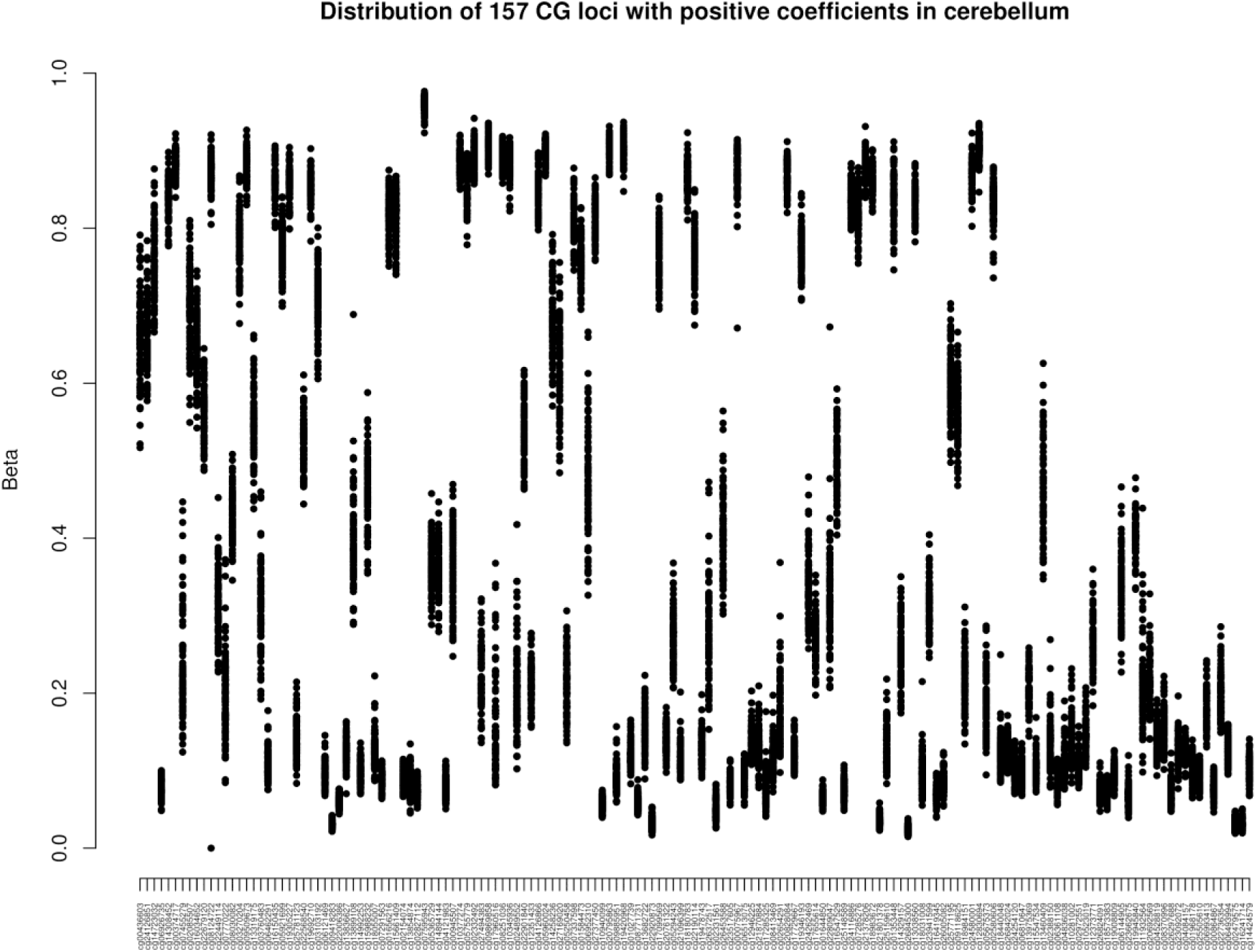
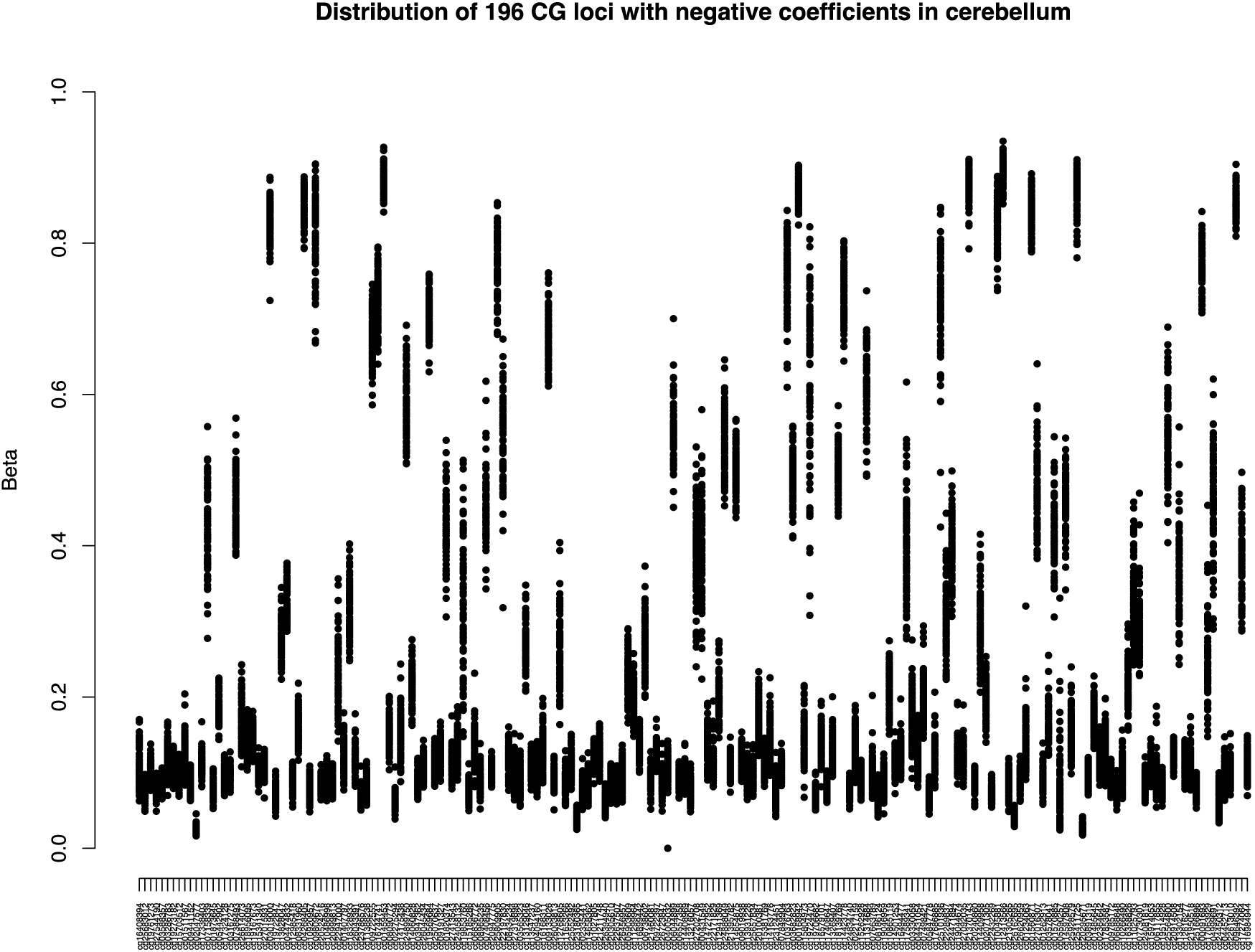
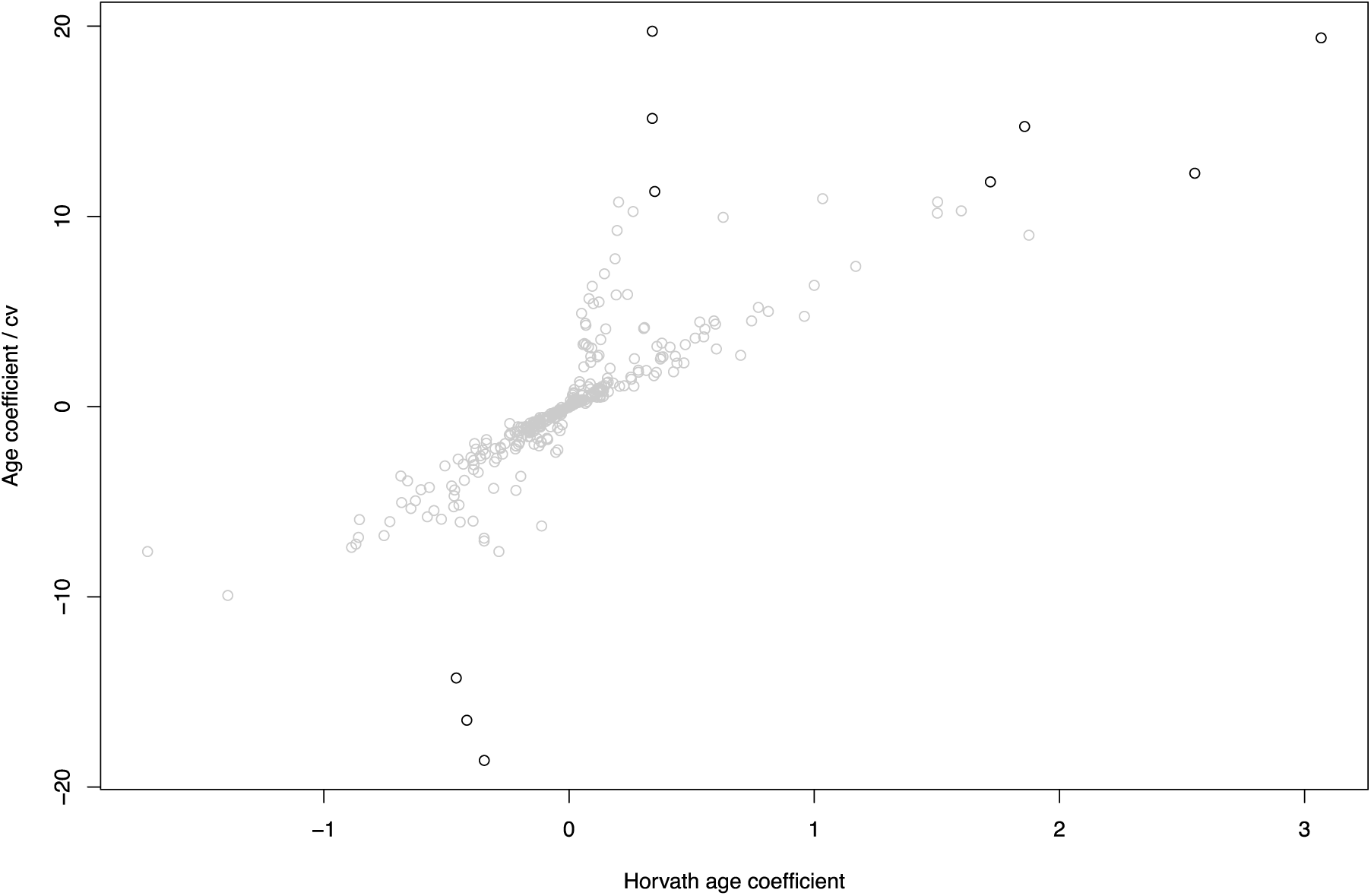
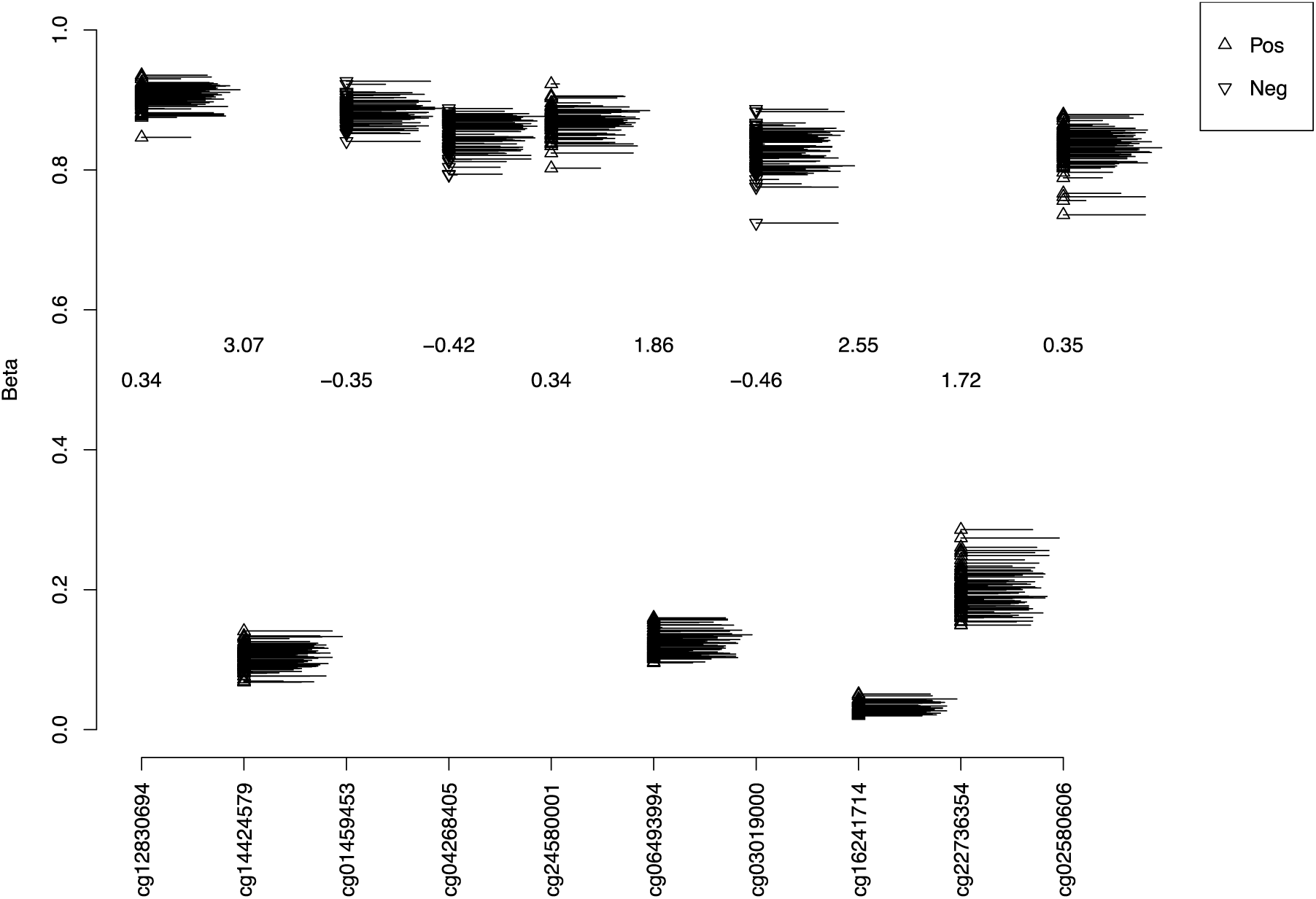
A) Exploration of Chronological vs DNAm ages and B) weighted average DNA methylation of the 353 Horvath clock CpGs vs chronological age in a dataset combining GSE40360, GSE53162, GSE59457, GSE61380, GSE61431, GSE67748, GSE67749, GSE89702, the London, and the Mount Sinai cohorts (Frontal lobe n=446, temporal lobe n=504, occipital lobe n=72, cerebellum n=246, blood n=80). An increase in divergence is observed as chronological age increases and a reduced slope among all samples (dotted curve) and especially the cerebellum samples (solid curve) can be noticed starting from the age of 60 which suggests the importance of introducing another inflection in the model at around that time. The solid line in fig 2a is the y=x bisector line.

The Horvath (2013) model includes 353 coefficients, but most of the magnitudes are clustered near 0 (figure 3a), leading us to hypothesize that the few largest coefficients might be particularly influential. In figure 3b, the distribution of DNA methylation proportions in cerebellum of the London cohort data is shown in a stripchart for each of the 157 loci with positive coefficients in the Horvath (2013) model, sorted by magnitude, with the largest coefficients at the right, and figure 3c for the negative coefficients (smallest at the left). It is striking that several loci with the largest positive and negative age coefficients in the model have low variance in this sample and also low DNA methylation.

**Figure 3.**
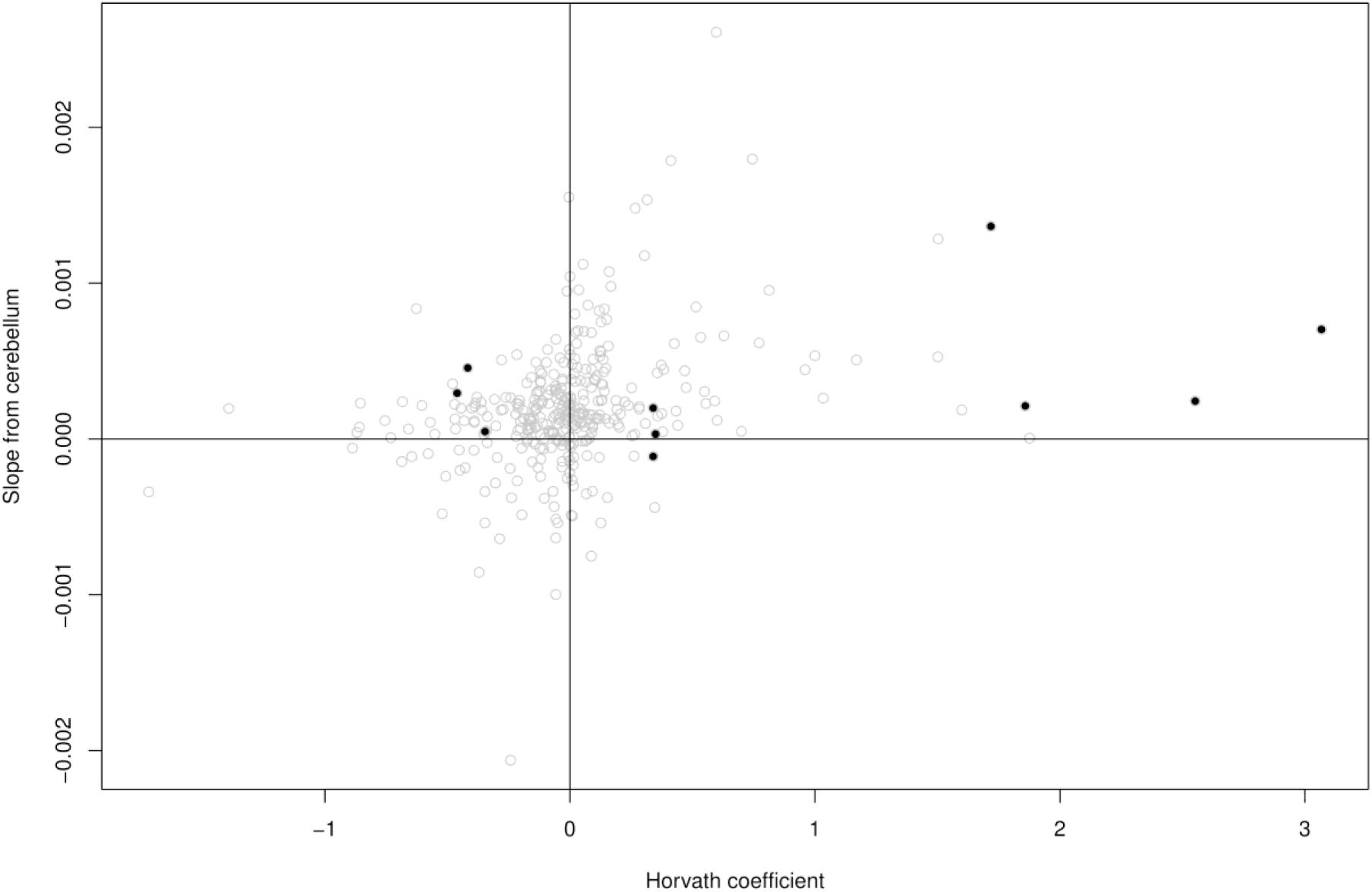
Exploration of model coefficients in the elderly cerebellum. a) stripchart showing the distribution of the 353 coefficients form Horvath (2013), b) stripchart showing the distribution of DNA methylation values of the 158 CG loci with positive coefficients in 83 cerebellum samples, c) stripcharts showing the distribution of DNA methylation values of the 195 CG loci with negative coefficients in 83 cerebellum samples, d) Scatter plot of age coefficients against their influence score (coefficient from Horvath (2013)/coefficient of variation in our data). The ten most influential loci are plotted in black, e) ten most influential loci, with the ages represented as a rug on the right hand side of each strip chart. The Horvath coefficients are shown in the center, and their sign is also denoted by the direction of the triangles, upward facing for positive and downward facing for negative, f) scatter plot of Horvath (2013) coefficients against their linear-model age coefficients in our data. The 10 most influential probes are shown in black.

One possible mechanism for the attenuated association of many CG sites in the model with age would be saturation (i.e. individual loci with positive coefficients reaching full methylation or negative coefficient ones reaching zero). To investigate this, we estimate the influence of each locus on the age estimate by dividing the coefficient from Horvath (2013) by an index of dispersion from our data, the coefficient of variation. In figure 3d we explore this value, and see that the most influential probes (black points at the top and bottom of the plot) include examples of both small variance (and large coefficient) and large variance (and small coefficient). This is further explored in figure 3e in which the ten loci with the highest influence are shown, with the ages equivalent to the length of each line protruding out of each circle and forming the rug on the right hand side of each strip chart. The rugs, scaled to best reflect the age range in the space available, give an indication of the noisiness of the relationship of the individual betas with age: the age relationship is not immediately visible for individual probes. The coefficients are shown in the centre, and the sign is also denoted by the direction of the stripchart, downward facing triangles for negative. This shows that of these ten probes there are three (cg12830694, cg24580001, and cg02580606) that might be candidates for saturation because they are highly methylated and expected to increase further with age. We thus fitted a regression line between chronological age and the beta values of each of the 353 loci and plotted the slopes against the Horvath coefficients. The 157 positively correlated and the 196 negatively correlated CpGs with age in Horvath (2013) are plotted and the ten most influential loci are highlighted in black (fig 3f). Of the ten most influential loci, four have a slope opposite in sign to the Horvath coefficient where three loci (cg08090772, cg03019000, and cg04268405) with negative Horvath coefficients have a positive slope in the cerebellum, and one locus (cg24580001) with a positive Horvath coefficient displays a negative slope in the cerebellum. In the main, the larger coefficients demonstrate the same direction of effect in our data as in the Horvath model. Admittedly this analysis is based on one tissue and age range, but it appears that many of the smaller coefficients may be effectively random with no biological meaning, and present in the model due to overfitting.

It is worth noting that the cerebellum is characterised by elevated levels of 5-hydroxymethylcytosine (5hmC) (Lunnon et al., 2016). We found that 31 out of the 353 Horvath clock sites were amongst the 65663 elevated 5hmC probes found in the cerebellum by Lunnon *et al* (2016). Of these, two sites (cg04268405, and cg24580001) are among the most influential sites reported in figure3e. Given that 5hmC is not distinguished from 5mC following bisulfite conversion, it is possible that age-associated changes to the 31 5hmC sites in the Horvath algorithm are offsetting the age predictions.

Finally, we examined whether age acceleration (calculated as the difference between DNAm age and chronological age) associates with AD neuropathology (measured using Braak score). Our results do show a weak association in some brain tissues (table 2). However, when age is included as a covariate, the association between age acceleration and AD pathology disappears (table 2). We also see this in the Mount Sinai cohort where no correlation was found between age acceleration and amyloid plaque levels in either the PFC or the STG when age is included as a covariate (table 2).

**Table 2.**
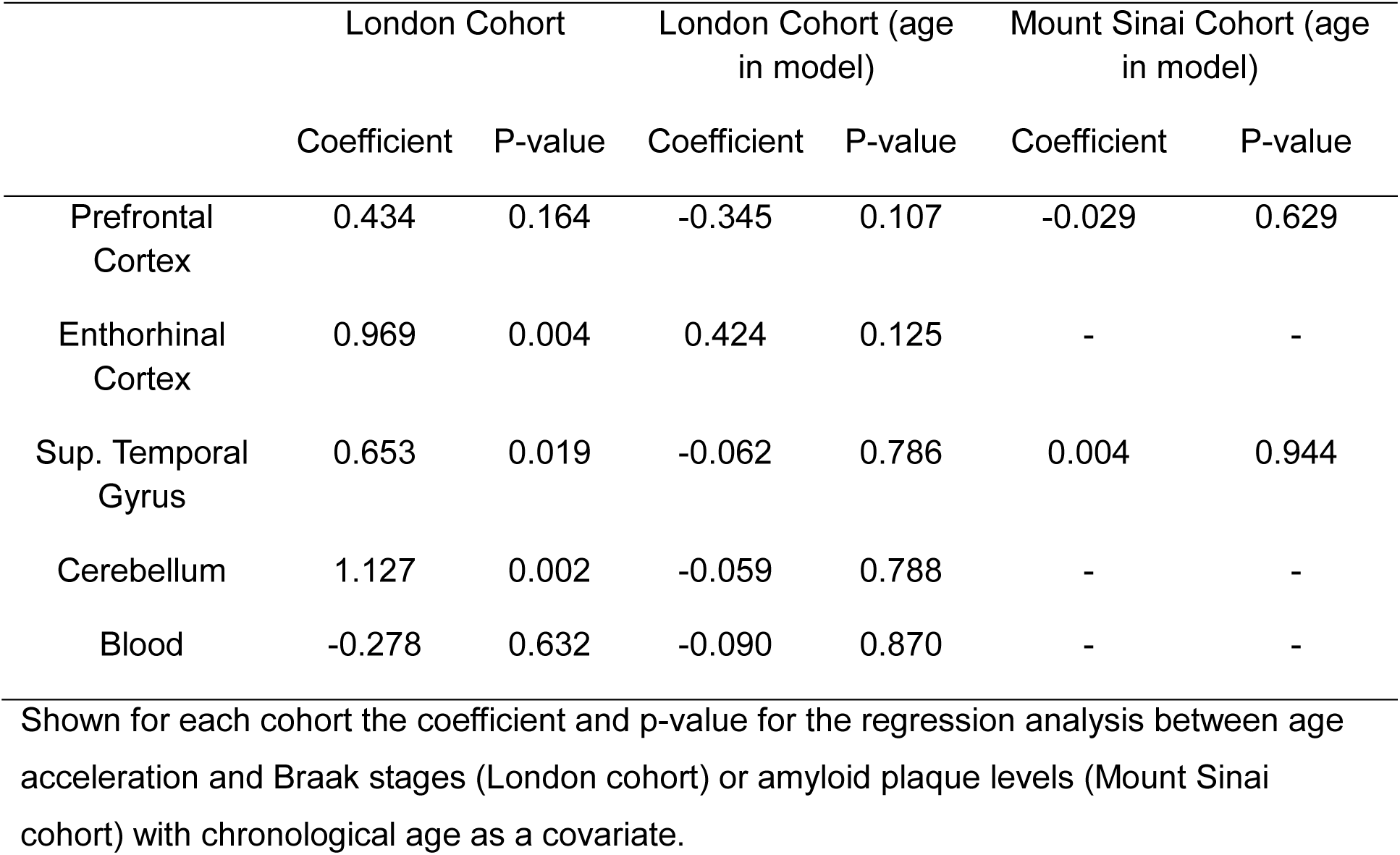
Regression analysis of epigenetic age acceleration of four brain tissues and blood from the London Brain Bank cohort versus brain Braak stage and of two brain tissues from the Mount Sinai cohort versus amyloid plaque levels. Both analyses were corrected for the chronological ages.

|  | London Cohort |  | London Cohort (age in model) |  | Mount Sinai Cohort (age in model) |  |
| --- | --- | --- | --- | --- | --- | --- |
|  | Coefficient | P-value | Coefficient | P-value | Coefficient | P-value |
| Prefrontal Cortex | 0.434 | 0.164 | -0.345 | 0.107 | -0.029 | 0.629 |
| Entorhinal Cortex | 0.969 | 0.004 | 0.424 | 0.125 | - | - |
| Sup. Temporal Gyrus | 0.653 | 0.019 | -0.062 | 0.786 | 0.004 | 0.944 |
| Cerebellum | 1.127 | 0.002 | -0.059 | 0.788 | - | - |
| Blood | -0.278 | 0.632 | -0.090 | 0.870 | - | - |

Shown for each cohort the coefficient and p-value for the regression analysis between age acceleration and Braak stages (London cohort) or amyloid plaque levels (Mount Sinai cohort) with chronological age as a covariate.

## Discussion

The Horvath (2013) ‘clock’ underestimates age in tissues and blood from older individuals. Following several publications listing loci at which DNA methylation associates with age (Hernandez *et al.*, 2011; Koch and Wagner, 2011), Horvath (2013) used penalized regression to train a multi-locus, cross tissue predictor of age using a large collection of public data. This involved a nonlinear transformation of the data to prevent the huge changes of childhood and adolescence from dominating the model. The resulting ‘epigenetic clock’ has been of practical use in predicting the age of unknown samples and as a quality check in epigenetic research. Additional widely used age predictors specific for blood were published by Hannum (2013) and Levine (2018) (phenotype-based). Here we analyse the Horvath (2013) clock, but the methods and many of the conclusions may be more widely applicable.

If the epigenetic clock is an index of an underlying ageing program that adapts to health and environment, then it resembles the circadian clock, and a biochemical mechanism for it remains to be discovered, particularly for adulthood and old age. Alternatively, it may represent the accumulation of DNA methylation change over time, analogous to a phylogenetic clock. When comparing DNA methylation profiles across tissues, individuals and other variables such as health, the dominant source of variation is the tissue, or more precisely the cell type. It is reasonable to suppose that this developmental blueprint can change over time in response to the environment, or simply drift or decay. This point of view corresponds roughly with the ‘Epigenetic Maintenance’ model posited by Horvath (2013).

Although the Horvath (2013) paper proposed an ‘epigenetic maintenance’ model of the epigenetic clock, which does not require an internal clock mechanism, the same work also featured the somewhat contradictory idea of ‘age acceleration’ in which discrepancies between DNA methylation (DNAm) age and chronological age might tell us something about a clock mechanism and the biological ageing status of the organism. These ideas are developed further by Horvath and Raj (2018). It is worth highlighting that in the latter third of the human age range, where you might most expect to see such associations, negative age acceleration increases with age. This means that any phenotype associated with age will appear to be associated with age acceleration as well, and a correct analysis should include chronological age as a covariate.

In a broad but non-comprehensive survey of the literature (table S1), we observe a variety of methods of calculating age acceleration, and many studies that do not correct for chronological age. Initially Δ-age (the difference between chronological age and the DNAm predicted age) was reported but alternative methods have since arisen: (1) the residual of regressing DNAm predicted age on chronological age (possibly in a model including covariates), (2) AgeAccel (difference between DNAm age value and the value predicted by a regression model in the control group), and (3) intrinsic (IEAA) and (4) extrinsic epigenetic age acceleration (EEAA) methods. Both IEAA and EEAA are methods applicable only on blood tissue since they subtract out the effect of blood cell count. One difference between the two, however, lies in that IEAA is based on the Horvath clock and does not take into consideration age-related decline of immune cells (Levine *et al.*, 2015). Contrarily, EEAA is defined on the basis of a weighted average of the epigenetic age from the Hannum clock and immune cells known to change with age (Horvath and Ritz, 2015).

In light of this methodological variety, we are concerned that the different epigenetic clocks, and the variety of age acceleration methods to choose from, would be laying a multiple testing trap that is overlooked by some colleagues in the field.

## Conclusion

Every adult experiences change over time in a way that makes the concept of ‘biological age’ compelling. The clock model has extremely interesting and useful characteristics but it is an extremely narrow summary of the DNA methylation profile. Many studies use age acceleration as the single epigenetic variable, based on only 353 CpG sites representing 1.15×10^−5^% of the methylome. Caution is urged when using the epigenetic clock, in particular in case control studies where studies should be carefully matched for age and may require further adjustment for age. Highly-powered epigenetic studies will continue to show some intriguing associations between age acceleration and varied health outcomes no matter which vision is true.

Finally, an ideal epigenetic clock and age predictor might be within reach but it would require the incorporation of all epigenetic changes observed with age and not just DNA methylation. Until then, it would be recommendable and more accurate to refer to the existing clocks as DNA methylation clocks instead.

## Methods

### Samples

#### Tissue Samples

Brain tissue samples (London cohort) were obtained from individuals diagnosed with Alzheimer’s disease (AD, n = 61) and from non-demented elderly control individuals (CON, n = 31) through the MRC London Neurodegenerative Disease Brain Bank as described in Lunnon *et al* (2014). In total four brain regions were analyzed (prefrontal cortex (PFC), the enthorhinal cortex (EC), the superior temporal gyrus (STG), and the cerebellum (CER)) and pre-mortem blood from a subset of individuals, collected as part of the Biomarkers of AD Neurodegeneration study. A second independent cohort (Mount Sinai cohort) was obtained from the Mount Sinai Alzheimer’s disease and Schizophrenia Brain Bank. This cohort consisted of two brain regions (PFC and STG) for 75 AD and 72 CON donors (Smith *et al.*, 2018).

#### Population sample: The UK Household Longitudinal study (UKHLS)

UKHLS is an annual household-based panel study which started collecting information about the social, economic and health status of its participants in 2009. Our analysis data set is drawn from one of the arms of UKHLS, namely, the British Household Panel Survey (BHPS), which merged with UKHLS in 2010 at the start of wave two. UKHLS collected additional health information, including blood samples for genetic and epigenetic analysis, at wave 3 for BHPS.

(www.understandingsociety.ac.uk)

#### Methylomic Profiling

DNA from the London cohort tissue samples were bisulfite-treated using Zymo EZ 96 DNA methylation kit (Zymo Research) according to the manufacturer protocol. DNA methylation levels were assessed on an Illumina HiScan System using the Illumina Infinium HumanMethylation450 BeadChip as previously described by Hannon *et al.* (2015). Raw signal intensities and probes for the London cohort were extracted using Illumina Genome Studio software were transformed into beta values using the Bioconductor wateRmelon package (Pidsley *et al.*, 2013). These were later normalized using the method implemented in the Horvath (2013) script. Data is available from both cohorts under GEO accession numbers GSE59685 (London cohort) and GSE80970 (Mount Sinai cohort).

1193 DNA samples from UKHLS were bisulfite-treated using Zymo EZ 96 DNA methylation kit (Zymo Research) according to the manufacturer protocol. DNA methylation levels were assessed on an Illumina HiScan System (Illumina) using the Illumina Infinium Epic Methylation BeadChip and samples were randomly assigned to chips and plates to minimise batch effects. Furthermore, in order to resolve any experimental inconsistencies, and to approve data quality, a fully methylated control (CpG Methylated HeLa Genomic DNA; New England BioLabs, MA, USA) was included in a random position on each plate. Raw signal intensities and probes for UKHLS were extracted using Illumina Genome Studio software and were transformed into beta values using the Bioconductor bigmelon package (10.18129/B9.bioc.bigmelon). These were later normalized using dasen function from the wateRmelon package (Pidsley *et al.*, 2013). After QC a final n of 1175 was reached.

#### DNA Methylation Age prediction

DNA methylation (DNAm) age was assessed for all samples of the London and Mt Sinai datasets on the the R statistical environment (R Development Core Team, 2015) using the script provided by Horvath (2013) as well as through the online DNAm Age Calculator (https://dnamage.genetics.ucla.edu/). These methods predicted age based on the DNAm coefficients of 353 CpG sites. The model (although not the custom normalization method) is also implemented in the agep() function of the wateRmelon package (version 1.17.0). This function was used to predict the ages of the UKHLS samples.

To assess the weighted average methylation levels, and to maximize the number of brain samples included in our assessment of age prediction, publically available 450KMethylation brain tissue datasets obtained from GEO (GSE40360, GSE53162, GSE59457, GSE61380, GSE61431, GSE67748, GSE67749, and GSE89702) along with the London and Mount Sinai cohorts were analysed (table 3).

**Table 3.**
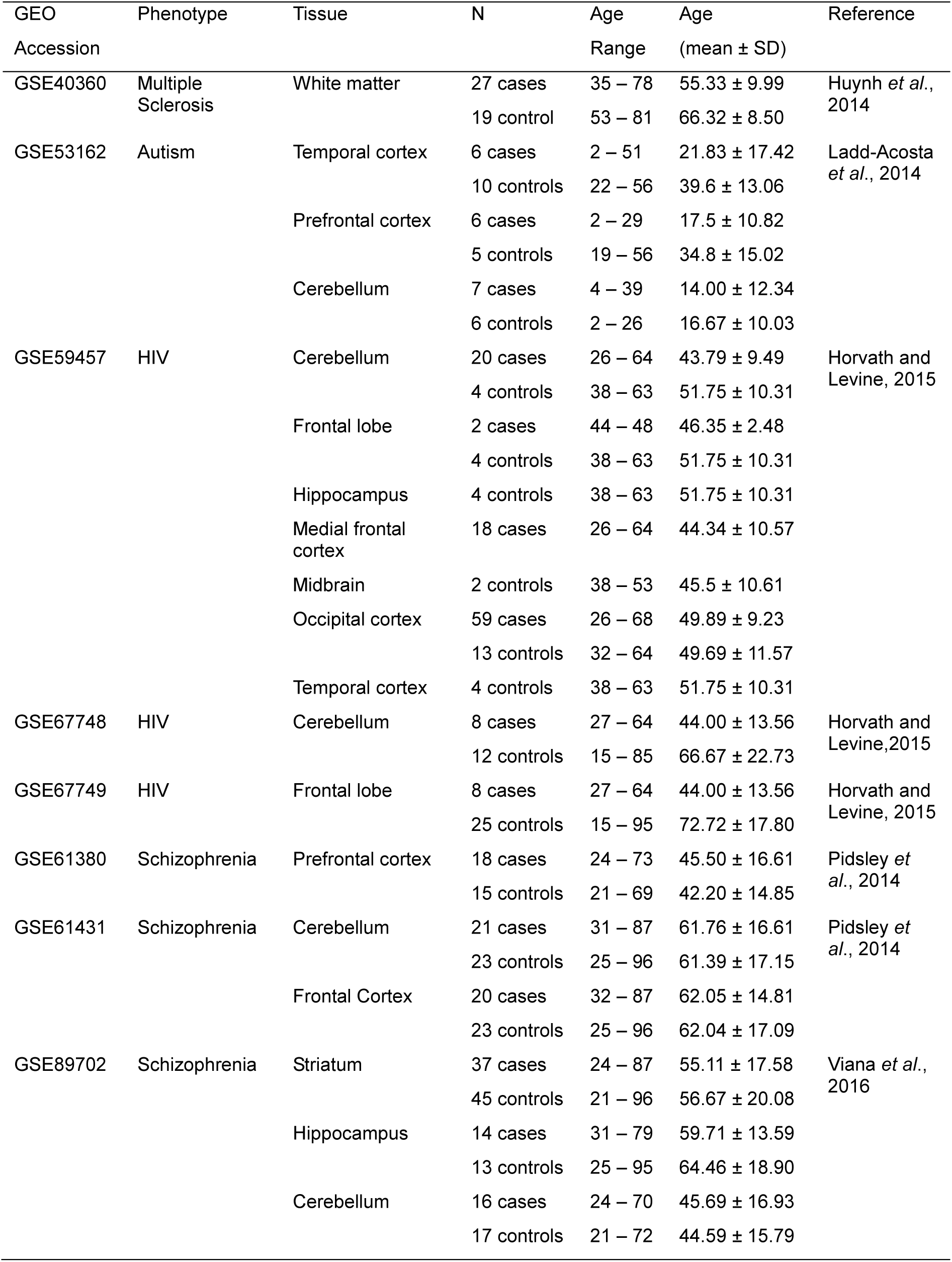
Breakdown of the phenotypes, tissues, and ages of the additional brain datasets

| GEO<br>Accession | Phenotype | Tissue | N | Age<br>Range | Age<br>(mean ± SD) | Reference |
| --- | --- | --- | --- | --- | --- | --- |
| GSE40360 | Multiple<br>Sclerosis | White matter | 27 cases<br>19 control | 35 – 78<br>53 – 81 | 55.33 ± 9.99<br>66.32 ± 8.50 | Huynh *et al.*,<br>2014 |
| GSE53162 | Autism | Temporal cortex | 6 cases<br>10 controls | 2 – 51<br>22 – 56 | 21.83 ± 17.42<br>39.6 ± 13.06 | Ladd-Acosta<br>*et al.*, 2014 |
| | | Prefrontal cortex | 6 cases<br>5 controls | 2 – 29<br>19 – 56 | 17.5 ± 10.82<br>34.8 ± 15.02 | |
| | | Cerebellum | 7 cases<br>6 controls | 4 – 39<br>2 – 26 | 14.00 ± 12.34<br>16.67 ± 10.03 | |
| GSE59457 | HIV | Cerebellum | 20 cases<br>4 controls | 26 – 64<br>38 – 63 | 43.79 ± 9.49<br>51.75 ± 10.31 | Horvath and<br>Levine, 2015 |
| | | Frontal lobe | 2 cases<br>4 controls | 44 – 48<br>38 – 63 | 46.35 ± 2.48<br>51.75 ± 10.31 | |
| | | Hippocampus | 4 controls | 38 – 63 | 51.75 ± 10.31 | |
| | | Medial frontal<br>cortex | 18 cases | 26 – 64 | 44.34 ± 10.57 | |
| | | Midbrain | 2 controls | 38 – 53 | 45.5 ± 10.61 | |
| | | Occipital cortex | 59 cases<br>13 controls | 26 – 68<br>32 – 64 | 49.89 ± 9.23<br>49.69 ± 11.57 | |
| | | Temporal cortex | 4 controls | 38 – 63 | 51.75 ± 10.31 | |
| GSE67748 | HIV | Cerebellum | 8 cases<br>12 controls | 27 – 64<br>15 – 85 | 44.00 ± 13.56<br>66.67 ± 22.73 | Horvath and<br>Levine, 2015 |
| GSE67749 | HIV | Frontal lobe | 8 cases<br>25 controls | 27 – 64<br>15 – 95 | 44.00 ± 13.56<br>72.72 ± 17.80 | Horvath and<br>Levine, 2015 |
| GSE61380 | Schizophrenia | Prefrontal cortex | 18 cases<br>15 controls | 24 – 73<br>21 – 69 | 45.50 ± 16.61<br>42.20 ± 14.85 | Pidsley *et al.*, 2014 |
| GSE61431 | Schizophrenia | Cerebellum | 21 cases<br>23 controls | 31 – 87<br>25 – 96 | 61.76 ± 16.61<br>61.39 ± 17.15 | Pidsley *et al.*, 2014 |
| | | Frontal Cortex | 20 cases<br>23 controls | 32 – 87<br>25 – 96 | 62.05 ± 14.81<br>62.04 ± 17.09 | |
| GSE89702 | Schizophrenia | Striatum | 37 cases<br>45 controls | 24 – 87<br>21 – 96 | 55.11 ± 17.58<br>56.67 ± 20.08 | Viana *et al.*,<br>2016 |
| | | Hippocampus | 14 cases<br>13 controls | 31 – 79<br>25 – 95 | 59.71 ± 13.59<br>64.46 ± 18.90 | |
| | | Cerebellum | 16 cases<br>17 controls | 24 – 70<br>21 – 72 | 45.69 ± 16.93<br>44.59 ± 15.79 | |

## Declarations

-Ethics approval and consent to participate: London cohort-Ethical approval NHS was provided by South East London REC 3; Mount Sinai Cohort-Ethical approval for the project was provided by the University of Exeter Medical School Research Ethics Committee under application number 14/02/041; UKHLS-Participants gave informed written consent for their blood to be taken and stored for future scientific analysis. The UKHLS has been approved by the University of Essex Ethics Committee and the nurse data collection by the National Research Ethics Service (10/H0604/2).

-Availability of data and material: The data used in this publication is previously published and available under GEO accession numbers given in table 3. Individual DNA methylation levels are available on application through the European Genome-phenome Archive under accession EGAS00001001232 (https://www.ebi.ac.uk/ega/home). Specific details can be found here (https://www.understandingsociety.ac.uk/about/health/data). Phenotypes linked to DNA methylation data are available through application to the METADAC (www.metadac.ac.uk)

-Competing interests: The authors declare no competing interests.

## Authors’ contributions

LS designed the study, LEK, TGS and LS did the analyses, MS AH YB AA JB EH and TGS generated and curated the UKHLS DNA methylation data, MK and JM advised and oversaw the work and all authors were involved in drafting and approving the manuscript.

## Acknowledgements

*Understanding Society:* Analysis was facilitated by access to the Genome high performance computing cluster at the University of Essex, School of Biological Sciences.

